# Phenotype Spectrum reflects Synergies among the Cell Architecture over Stages of the Cell Cycle

**DOI:** 10.1101/2020.06.05.135723

**Authors:** Tian Lan, Meng Yu, Weisheng Chen, Jun Yin, Hsiang-Tsun Chang, Shan Tang, Ye Zhao, Spyros Svoronos, Samuel W. K. Wong, Yiider Tseng

## Abstract

The heterogeneity of cell phenotypes remains a barrier in progressing cell research and a challenge in conquering cancer-related drug resistance. Cell morphology, the most direct property of cell phenotype, evolves along the progression of the cell cycle; meanwhile, cell motility, the dynamic property of cell phenotype, also alters over the cell cycle. However, a quantifiable research understanding the strict relationship between the cell cycle and cell migration is missing. Herein, we separately elucidate the correspondence of single NIH 3T3 fibroblast migratory behaviors with the G1, S, and G2 phases of the cell cycle, an underlying property of proliferation. The results show that synergies among the highly spatiotemporal arrangements of signals in Rho GTPases and cyclin-dependent kinase inhibitors, p21^Cip1^, and p27^Kip1^ coordinates proliferation and migration. Taken together, we explain the synergies among these processes through providing an interactive molecular mechanism between the cell cycle and cell migration and demonstrate that both cell morphology and the dynamic subcellular behavior are homogenous within each stage of the cell cycle phases, posing potential implications in countering drug resistance.

## Introduction

Cell migration and proliferation both play essential roles in physiological and pathological events, including embryo development, wound repairs, and cancer metastasis. During vertebrate morphogenesis, cell populations continuously proliferate before migrating to desired locations to form an appropriate tissue pattern (Locascio & Nieto, 2001). In cancer metastasis, cells grow, extravasate, and eventually migrate into remote tissues, where they proliferate again to establish secondary tumors (Friedl & Wolf, 2003). Although proliferation and migration are two pertinent cellular activities that commonly coexist, it is poorly understood whether this coexistence is independent of each other or a result of interactive molecular mechanisms connecting the cellular activities.

Cell proliferation is funneled by the cell cycle, which is regulated by the cyclin-dependent kinases (CDKs) (Murray *et al*, 1989; Serrano *et al*, 1993; van den Heuvel & Harlow, 1993). When CDKs are activated, they trigger gene expression waves (Uhlitz *et al*, 2017) that have obvious impacts on the cell behavior. Hence, on a more rudimentary level, it is widely accepted that the alternation of cell morphologies and dynamics is also a natural consequence of the cell cycle progression. Despite observations surrounding these processes, the cell cycle’s impact on cell dynamics has not been systematically addressed before. Cell migration requires cytoskeletal remodeling, for which Rho GTPases serve as the primary molecular switches (Narumiya, 1996; Nobes & Hall, 1995). When RhoA, one of the Rho GTPases, is activated, it propagates its signal to downstream proteins, Rho Kinase (ROCK), then LIM kinase (LIMK) to inhibit ADF/cofilin in its F-actin severing and G-actin sequestering activities(Ohashi *et al*, 2000). The crosstalk between the CDKs and RhoA pathways that relates the cell cycle and cell migration have only been mentioned through two CDK inhibitors (CDKIs), p27^Kip1^ and p21^Cip1^ (Besson *et al*, 2004; Gui *et al*, 2014; Lee & Helfman, 2004; Phillips *et al*, 2018).

As the cell is quiescent in the G1 phase, p27^Kip1^ and p21^Cip1^ tightly bind to the cyclin-CDK complexes in the nucleus to prevent the cells from entering the cell cycle (Harper *et al*, 1993; Polyak *et al*, 1994). Upon mitogen stimulation, cells then enter the S phase as these CDKIs dissociate from the cyclin-CDK complexes so that the chromosome can be duplicated. Consequently, p27^Kip1^ is exported from the nucleus to the cytoplasmic region to bind and inhibit RhoA before the p27^Kip1^-RhoA complex is degraded (Besson *et al.*, 2004; Phillips *et al.*, 2018). On the other hand, p21^Cip1^ will bind to and inhibit the ROCK (Lee & Helfman, 2004). Since the RhoA signaling is disrupted by these two CDKIs after the G1/S phase transition, the reactivation of ADF/cofilin should allow actin-cytoskeletal remodeling (Arber *et al*, 1998; Carlier *et al*, 1997; Ohashi *et al.*, 2000). As a result, it has been speculated that CDKIs play a dual role in tumorigenesis to release cells from cell cycle arrest and in metastasis to enhance actin-cytoskeletal dynamics (Besson *et al*, 2008; Gui *et al.*, 2014; Lee & Kim, 2009; Phillips *et al.*, 2018; Rampioni Vinciguerra *et al*, 2019).

This perspective suggests that the interactions among CDKIs and proteins in the RhoA pathway after the G1/S phase transition increase the likelihood for the metastasis of cancer cells whereas cell spread events likely occur in the G2 phase. However, previous studies have reported that cell motility in the G1/S phases is greater than in the G2/M phases for all the cell types they surveyed, including L929, HeLa, and BT4Cn glioma cells (Walmod *et al*, 2004). Additionally, NBT-II rat bladder carcinoma cells respond to Fibroblast Growth Factor-1 and increase their dynamics only in the G1/S phases but not in the G2 phase (Bonneton *et al*, 1999). These cases reveal that the surveyed cancer cells only have their motility reduced after passing the G1/S phase transition and are against the proposal for the roles of these two CDKIs in cell dynamics. The gap in understanding whether cell motility should be used in evaluating cell invasion or whether we lack an integrative view for understanding cell dynamics in the G2 cells makes it critical to provide a robust framework for the relationship between the cell cycle and cell migration.

Hence, in this study, we apply a robust cell migration assessment, the *CN correlation* analysis (Lan *et al*, 2016; Lan *et al*, 2018) to probe the phenotype spectrum (*i.e.*, the alternation of cell morphologies and dynamics regarding the stages of the cell cycle). Rather than utilizing traditional cell migration analysis techniques, which obligate a long period of cell trajectories (~10 hours) (Noever *et al*, 1994; Wu *et al*, 2009) and provides a dilemma where the duration of the sampled cell trajectories must be long enough for statistically meaningful analysis, yet short enough to have the trajectories restrained within the same stage of the cell cycle phases, the *CN correlation* analysis allows us to decipher the migratory phenotype of cells in each stages of the cell cycle and to clarify the impact of the two aforementioned CDKIs on cell dynamics, hence obtaining the comprehensive relationship between cell migration and the cell cycle. Building this framework provides an architecture through which we can map the structural properties throughout the cell cycle and simplifies our understanding of cell physiology.

## Results

### The evolutions of both cell velocity and morphology stem from cell cycle progression

We first investigated whether the cell migration pattern aligns with cell cycle progression. To do this, we monitored the cell trajectory and the corresponding cell shape of an NIH 3T3 fibroblast throughout the cell cycle (**Fig. 1A**). This result revealed that the cell migratory trajectory is a straight line in the beginning but become curved later. Accordingly, the cell shape also transforms significantly. Hence, we scrutinized motility and polarity of the cell at 3-minute intervals over the whole cell cycle in terms of four basic cell parameters: instantaneous speed, area, aspect ratio, and circularity (**Fig. 1B**). We also applied the least-square criterion to classify the complete cell cycle into different periods based on the parameter similarities. The results suggest that the whole cell cycle could be roughly divided into three distinct periods (see Materials and Methods).

**Figure 1.**
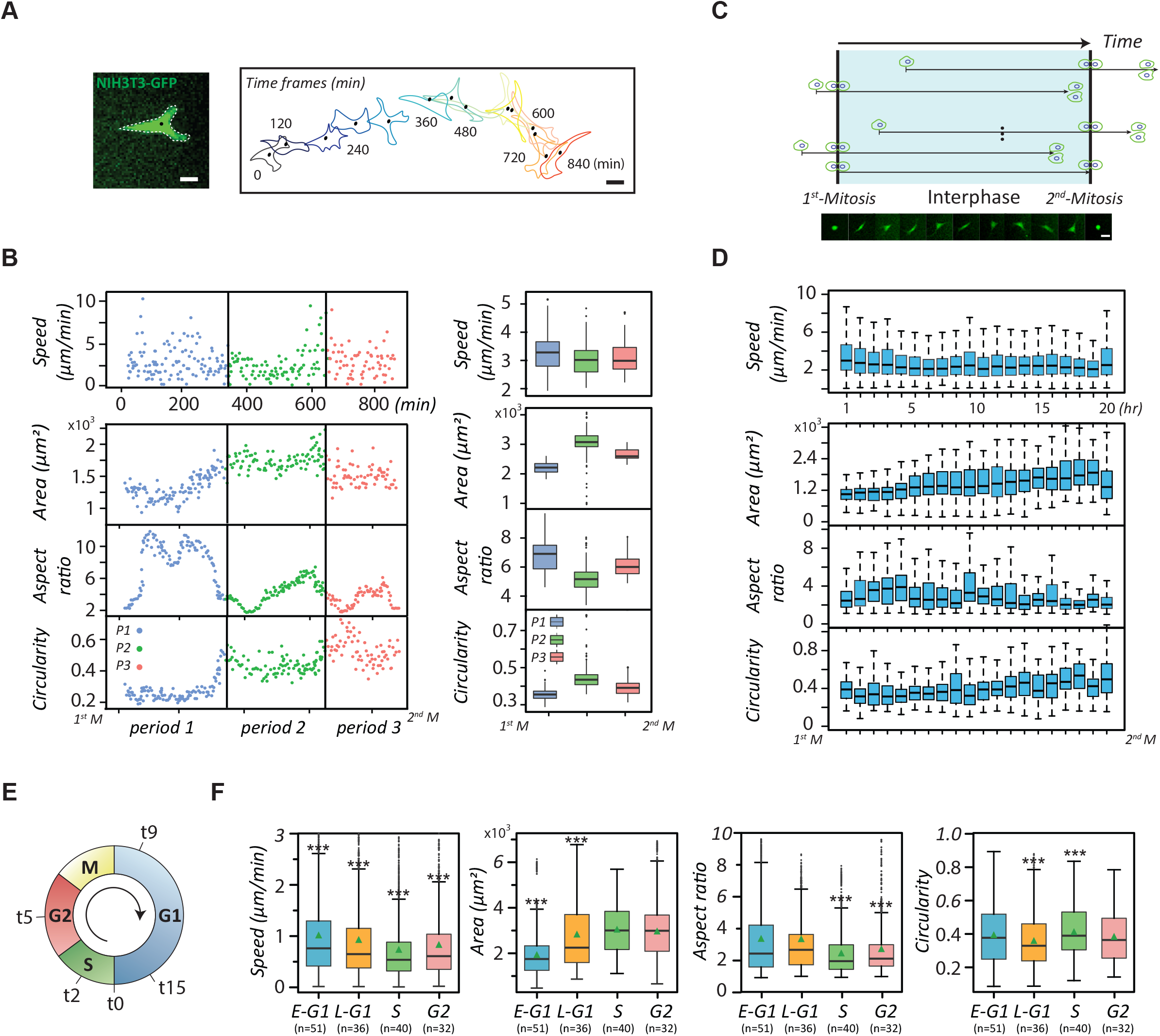
The evolution of migration and morphology of NIH fibroblasts over the cell cycle. (**A**) Green fluorescent protein (GFP)-labelled single NIH3T3 fibroblasts monitored at 3-min time intervals over an entire cell cycle beginning at cell-division. The time-evolved cell outlines and corresponding cell centroids over the entire cell cycle of a fibroblast are shown. Scale bar: 20 μm. (**B**) Scatter plots (left) and Box plots (right) showing the dynamic and morphological evolutions of single NIH 3T3 fibroblasts during the cell cycle progression. The cell phenotypes, including instantaneous speed, area, aspect ratio, and circularity were monitored at 3-minute time intervals throughout the cell cycle. The perpendicular lines separate periods of time exhibiting different cell phenotypes. (**C**) The schematic of the time course alignments of live, single cells tracked over 20 hours, related to Fig. 1D. The series of micrographs show the morphological evolution of the GFP-labeled NIH 3T3 fibroblasts over the whole cell cycle. Scale bar: 20 μm. (**D**) The box plots verifying cell phenotype evolutions over the cell cycle using 20 cells. (**E**) The plot representing the time of the optimal profiles of double-thymidine-synchronized cells that propagate to the early G1, the late G1, the S, and the G2 phase, determined as 9, 15, 2 and 5 hours, respectively (and denoted as t9, t15, t2, and t5, respectively). (**F**) The box plots showing the dynamic and morphological measurements of synchronized NIH 3T3 fibroblasts (n > 20) in different stages of the cell cycle phases. Data information: *** indicates that the group of data is statistically significant compared to other groups. The statistical results of this figure are listed in Table EV1.

Our results show that the cell’s instantaneous speed fluctuates abruptly in the first and third periods but maintains consistently low in the second period. The cell area is initially small but continuously increases in the first period, steadies in the second period and disperses vigorously in the third period. In terms of cell shape, the monitored cell displays a high aspect ratio and low circularity in the first period but contrasting results in the other periods, demonstrating that the cell shape transforms after the second period.

To verify that these results do not emerge from coincidence, we acquired 20 single-fibroblast movies, 20 hours long each, ensuring inclusion of at least one cell division event. We then identified a movie containing a complete cell cycle (*i.e.*, possessing two cell divisions) and set its timeframe as a benchmark to align the timeframes of the rest. If the cell division occurred at an earlier time in the aligned movie, then the division time of this movie would be superimposed with the first division of the standard one; otherwise, the division time would be overlaid with the second division of the standard one (**Fig. 1C**). Under these conditions, all cells possessed relatively similar timeframes with respect to the cell cycle. Hence, we could analyze cell phenotypes of the cells appearing in those movies over the reference timeframe using the four-cell parameters (**Fig. 1D**). The results from this assessment confirm the existence of three distinct periods.

The presence of three distinct phenotypes within the cell cycle seems to correlate with the progression of the cell cycle. Therefore, we further synchronized NIH 3T3 fibroblasts into different stages of the cell cycle phases (**Fig. 1E** and **Fig. EV1**) and randomly acquired 25 single-cell movies in each of these stages to re-analyze these parameters (**Fig. 1F**). The results show that the instantaneous speeds are the highest in the G1 phase, reach the minimum in the S phase, and slightly bounce back in the G2 phase. The cell area displays a monotonically increasing trend until the late G1 phase and reaches a plateau after entering the S phase. The aspect ratio and circularity together suggest that cells have the lowest polarity in the S phase. In essence, cell migration exhibits a strong correlation with the cell cycle.

### Overview: *CN-correlation* assessment

To probe cells’ migration properties in each cell cycle phase, we developed a novel biological model, termed the *CN correlation analysis*, to statistically decode cell motions (Lan *et al.*, 2016). Briefly, we composed a *CCD vs. NCD*_//_ coordinate system (the *CN* plot), where *CCD* is the cell centroid displacement and the *NCD*_//_ is the relative nuclear centroid displacement. Then, the *CN correlation* data were collected from single-cell movies, acquired at one-minute intervals, with individual datum being deciphered from two adjacent movie frames (**Fig. 2A**).

**Figure 2.**
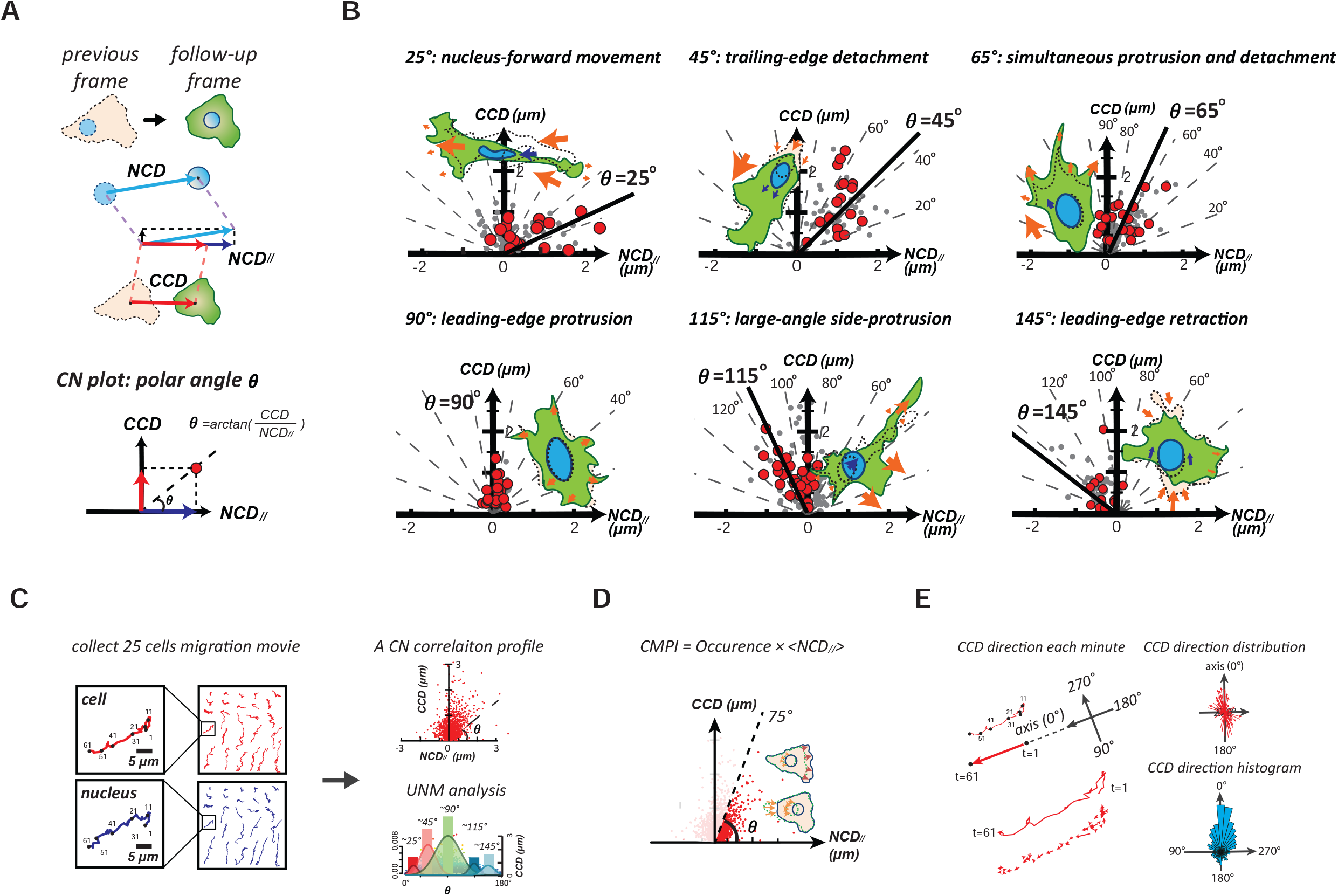
The overview of CN correlation analysis. (**A**) The framework of *CN correlation* analysis. (**B**) Six representative subcellular activities displaying their specific distributions in the *CN* plot. A signature *CN polar angle* represents the migratory pattern of the corresponding subcellular activity. (**C**) Illustration of the *CN* profile definition and the univariate normal mixtures (*UNM*) analysis. A *CN correlation* profile (n = 1500) is constructed using a collection of 25 randomly selected, one-hour movies of NIH fibroblasts at one-minute intervals. The *UNM* analysis is applied to profile the composition of normal distribution curves that describe the proportion and signature migration patterns. (**D**) The schematic of the cell migration potential index (*CMPI*) according to the equation: *CMPI* = *Occurance* ×< *NCD*_//_ >. Occurrence denotes the frequency rate of the *CN correlation*s over populations located in the region (0°-75°); <*NCD*_//_> represents the mean value of *NCD*_//_ over the whole population. (**E**) Methodology schematic for the migration polarity analysis of the cell. One-minute *CCD* direction is aligned with the orientation of one-hour displacement. The direction histogram reveals migratory polarity.

Accordingly, we obtained a series of *CN correlation* data from an identified cell locomotion event in cell movies and evaluated their distributions in the *CN* plot. We found that data generated from each specific locomotion event follow a specific polar-angle-distribution pattern: The peak polar angle of nucleus-forward movement is ~25°, trailing-edge detachment is ~45°, simultaneous protrusion and detachment is ~65°, leading-edge protrusion is ~90°, large-angle side-protrusion is ~115°, and leading-edge retraction is ~145° (**Fig. 2B**). We denote these angles as signature polar angles, and the *CN* plot with data as the *CN* profile.

Conversely, the *CN correlation* analysis can also be applied to an undesignated cell movement event to reveal the percentages of individual subcellular activities contributing to the event. When the polar angle profile of the *CN* plot from a probed cell movement event is subjected to the automatic univariate normal mixtures (*UNM*) algorithm to statistically decompose the profile into distinct normal distributions, the peak values of which will separately map to the signature polar angles of the corresponding subcellular activity and the size of the normal distribution will reveal the weight of the subcellular activity in the migration pattern (**Fig. 2C**). Hence, the *CN* profile is a statistical descriptor that analyzes the unique cell migration pattern of a specific cell type. This information can also be further extended to the underlying signaling pathways involved in each subcellular activity (Lan *et al*, 2018).

Since cell motility is carried out by effective migration, a cell and its coupled nucleus must move in the same direction. In a *CN* profile, only the data located within the polar angles ranging from 0° to 75° are considered to contribute to the effective migration. Hence, cell motility of a cell type under a certain condition is estimated by the sum of the effective nuclear displacements in the *CN* profile, called Cell Migration Potential Index (*CMPI*) (**Fig. 2D**). Greater *CMPI* values of a cell indicates faster motility over a long period of time. Our previous study had shown that the *CN correlation* analysis possesses excellent applicability and reliability for cell migration assessment (Lan *et al.*, 2016; Lan *et al.*, 2018), so we incorporated cell polarity into the *CN correlation* analysis to evaluate the persistence of mesenchymal cell migration. The consistency of the short-term *CCD* orientations with respect to the long-term cell displacement indicates that the probed cell possesses high polarity (**Fig. 2E**). Hence, the *CN correlation* analysis can assess cell migration from both the motility and polarity perspectives.

### Cell migration patterns are distinct and highly homogeneous in each cell cycle phase

We first assessed the different migration patterns appearing in different stage of the cell cycle through cell morphology changes. We compared the cell images of the same cell acquired under 3-minute intervals at every 10° of cell boundaries to characterize the changes in cell morphology, as shown in the representing cells (**Fig. 3A**). The results reveal that cells display distinct migration patterns in each stage of the cell cycle. Cells in the G1 phase are highly motile, cells in the S phase are relatively still, and cells in the G2 phase exhibit notable side-protrusion. Interestingly, in the early G1 phase, cell motions follow along a fixed axis (*i.e.*, in a consistent direction) but this trend gradually diminishes while approaching the S phase. When cell motions resurge in the G2 phase cells, these motions do not align at a fixed axis.

**Figure 3.**
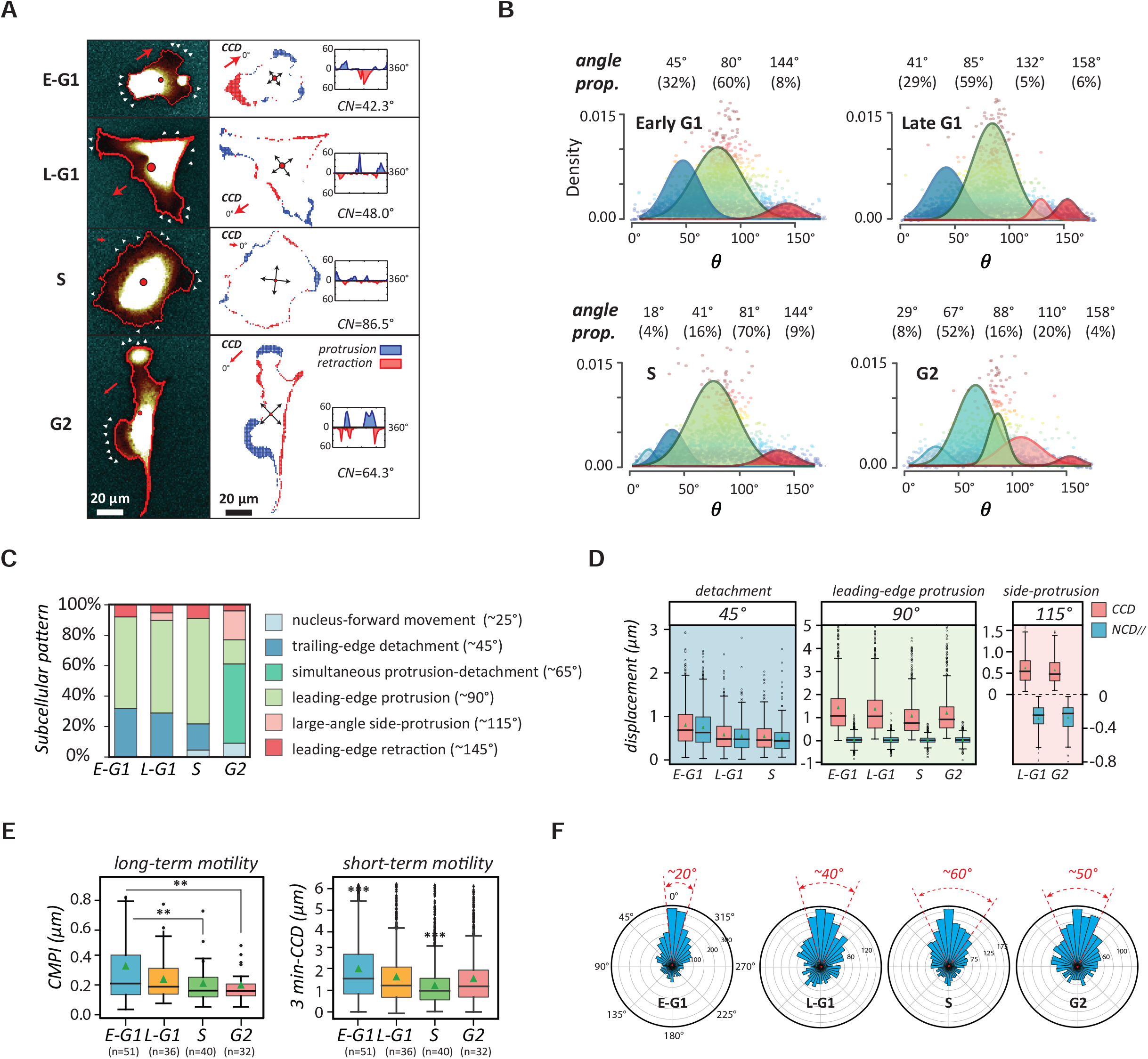
The transition of natural migration patterns of NIH 3T3 fibroblasts over the cell cycle. (**A**) Graphic of typical spreading patterns of NIH 3T3 fibroblasts in each stage of the cell cycle phases. Cell migratory dynamics can be assessed by peripheral variations in the polar coordinate in which cell centroid is set as the origin – *left column:* fluorescent cell images in different cell cycle phases; *right:* the occurrences of protrusions (blue) and detachments (red) in the corresponding synchronized cell edges; *insert:* the distribution of protrusion and retraction. (**B**) The decomposed *UNM* signature plots of synchronized cells showing cell migration patterns in the different cell cycle stages, where the peak polar angle of each *CN correlation* data distribution represents a distinct subcellular activity with specific weights. (**C**) Cell migration patterns of NIH 3T3 fibroblasts, decomposed into different subcellular events. (**D**) Box plots of CCD and NCD_//_ in each signature polar angle zones, representing the magnitude of distinct subcellular events. (**E**) Cell motilities in different stages of the cell cycle phases, estimated using the Cell Migration Potential Index (*CMPI*) for the long-term and three-minute *CCD* speed for the short-term. (**F**) Cell migration polarity in different stages of the cell cycle, evaluated by the one-minute *CCD* direction histogram. Data information: *** indicates that the group of data is statistically significant compared to other groups. The related statistical results of this figure are listed in Table EV2.

Hence, we applied the *CN correlation* analysis to systematically characterize the migration patterns of single NIH 3T3 fibroblasts, synchronized within each cell cycle phase (**Fig. 3B** & **Table 1**). This analysis not only specifies unique motion patterns in distinct stages of the cell cycle, but also illustrates the evolution of phenotypical transition of cells through the cell cycle. Moreover, since the results are highly resolved, they also verified the homogeneity of cell migration patterns within each stage of the cell cycle (**Fig. 3C**).

**Table 1.** The proportions of subcellular activities contributed to the cell migration patterns of cells in different cell cycle phases. The table listing the subcellular activities with signature polar angles and their percentiles in each stages of the cell cycle phases, determined by the *CN correlation* analysis.

| The cell cycle phase | Early G1 | Late G1 | S | G2 |
| --- | --- | --- | --- | --- |
| ~25°: nucleus-forward movement | - | - | 18° (4%) | 29° (8%) |
| ~45°: trailing-edge detachment | 45° (32%) | 41° (29%) | 41° (16%) |  |
| ~65°: simultaneous detachment-protrusion | - | - |  | 67° (52%) |
| ~90°: leading-edge protrusion | 80° (60%) | 85° (59%) | 81° (70%) | 88° (16%) |
| ~115°: side-protrusion | - | 132° (5%) | - | 110° (20%) |
| ~145°: leading-edge retraction | 144° (8%) | 158° (6%) | 144° (9%) | 158° (4%) |

In the early G1 phase, we found that cells exhibit a standard, highly polarized fibroblast behavior, where detachment events take ~30% of the time and protrusion events take ~60%. In the late G1 phase, the cells deviate from the initial directional migratory mode and their polarity fades, judging by the increase of side protrusion events and the reduction of *CCD* and *NCD*_//_ (**Fig. 3D**). In the S phase, the detachment events drop significantly to 16%, while the protrusion events are dominant at 70%. Meanwhile, the sizes of *CCD* and *NCD_//_* are both at a minimum (Fig. 3D), indicating the cells have lost their polarities and intend to remain stationary. Notably, upon reaching the G2 phase, the cells only perform pure protrusion 16% of the time. In contrast, 52% of the time the cells are predominantly occupied by simultaneous detachments and protrusions. In addition, the cells also exhibit large-angle side protrusion 20% of the time. This irregular dynamic pattern is well-illustrated by the G2-cells shown in the Fig. 3A, where the cell displays neither a polarized nor an isotropic, but rather, an irregular shape.

Notably, *CMPI* analysis suggests that the motility of the cells in the G2 phase is low, comparable to in the S phase; however, the instantaneous speeds (3-minutes *CCD*) of the cells in the G2 phase are relatively high and similar to those in the late G1 phase (**Fig. 3E**). Hence, the high instantaneous speeds are not translated to the long-term motility for the G2-phase cells, a typical phenomenon for poorly polarized phenotypes. This conclusion is also further supported by the persistency evaluation (**Fig. 3F**), which shows that the *CCD* directions alters more frequently for the cells in the G2 phase than in the G1 phase.

### The actin-cytoskeletal remodeling leads to the cell-cycle-dependent migration pattern

Cell dynamics are controlled by cytoskeletal remodeling. Migration-related subcellular activities, including protrusion, detachment, contraction, and adhesion, all stem from spatiotemporally assembling and disassembling of actin filaments (Kirfel *et al*, 2004; Lauffenburger & Horwitz, 1996). Hence, we examined the structures of the actin-cytoskeleton and cell adhesions in different stages of the cell cycle through fluorescence microscopy (**Fig. 4A**). The representing micrographs show that the early G1-phase cells have an amorphous actin-network and sparsely distributed adhesion plaques, which hallmarks the motile pattern (Bohnet *et al*, 2006; Le Clainche & Carlier, 2008). In contrast, in the S and the G2 phase, cells form defined cytoskeleton with highly organized stress fibers and dense-patched cell adhesions, signifying a stable, immotile pattern (Kanchanawong *et al*, 2010).

**Figure 4.**
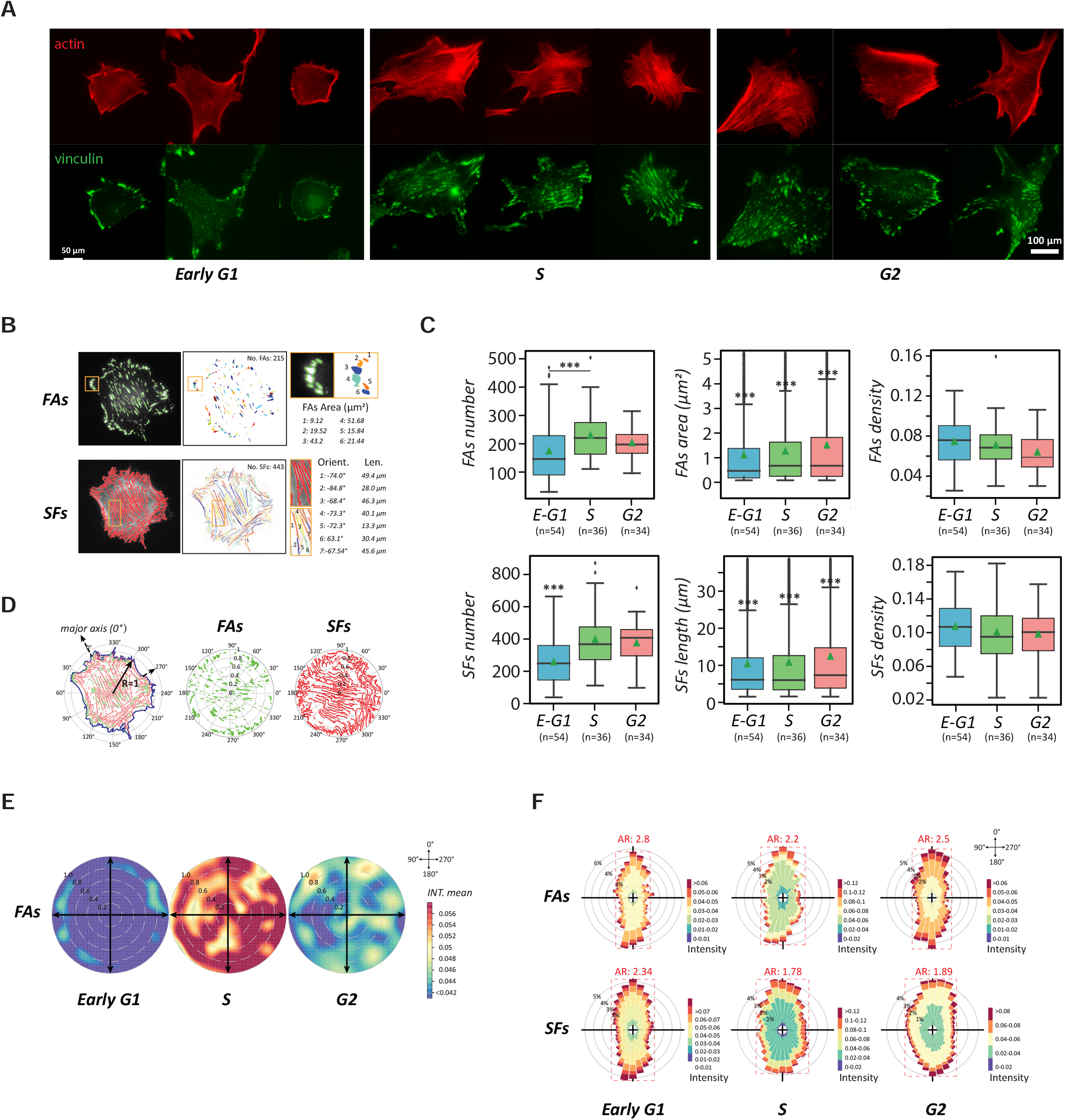
The transition of cell motility and polarity through the cytoskeletal remodeling over the cell cycle. (**A**) The cytoskeleton (actin; red) and adhesion plaques (vinculin; green) in NIH 3T3 fibroblasts over interphase of the cell cycle. Scale bar: 100 μm. (**B**) Segmented focal adhesions (FAs) and stress fibers (SFs) subjected to quantitative analysis using computer vision– *left column:* visualization of boundary segmentation of fluorescent cell images; *middle:* quantity of segmented FAs and SFs in single cells, shown in different color; *right:* the quantification of FA and SF features, including size, length, and orientation. (**C**) Box plots presenting FAs and SFs dynamics during the cell cycle progression using the total number, single FAs area/SFs length and density over the cell body. (**D**) The localization distribution of FAs in the cell normalized to a circle, where the distance to the centroid and the closest edge counts serve as the normalized position and the distribution is quantified by polar coordinates, in which cell centroid is set as the origin, and the 0° is aligned to the major axis. (**E**) Rose plots showing the distribution of FAs in each cell cycle stage at the cell population level. The degree of mean intensity is represented by warm (active)-cool (non-active) colormap. (**F**) Polar histogram plots showing the frequency localization of FAs (up) and SFs (down) of each cell cycle stage of the cell cycle phases, where AR denotes the aspect ratio of the frequency distribution shape. The greater value of AR represents a more polarized FAs/SFs distribution. Data information: *** indicated that the group of data is statistically significant compared to other groups. The related statistical results of this figure are listed in Table EV3.

To further understand the cell-cycle-dependent migration patterns, we scrutinized the stress fibers (SFs) and focal adhesions (FAs) in detail. The number, size (area), and density of the SFs and FAs were measured from the corresponding cell images (**Fig. 4B**), providing insightful cytoskeletal evolution throughout the cell cycle progression (**Fig. 4C**). In the early G1 phase, small adhesion plaques are scattered over the whole cell bodies. These adhesion plaques are not matured and could be disassembled easily; thus, the cells are highly motile (Choi *et al*, 2008). In the S phase, the overall numbers and sizes of FAs and SFs in a cell both increase, but their densities drop over the enlarged cell body, indicating that the adhesion plaques were merged into matured patches and the extended SFs became stable, which led to immotile cells. Even though the G2-phase cells continuously possess thick SFs and matured FAs from the S phase, the density of FAs are significantly reduced. This change inspires the speculation for the occurrence of focal adhesion turnover.

To clarify whether focal adhesion turnover had occurred, we applied polar coordinates to individual cells to normalize their cell shape and size dependence and statistically characterize the FAs within the cells (**Fig. 4D**). In the polar coordinate system, we set the origin at each cell’s centroid, the major axis of the cell in the direction of the leading edge as polar angle 0° for each cell, and the radii as the normalized distances from the cell centroid to the radiated cell edges. Following this format, FA distributions in different cells were overlaid on a rose plot with a colormap representing the density (**Fig. 4E**). The rose plots clearly demonstrate that cells in the S phase possess the highest density of adhesions. Therefore, the comparison of local adhesion densities between cells in the S and the G2 phase supports the idea that the FAs diminish in certain regions of the G2 cells and focal adhesion turnover occurs. As a result, parts of cell periphery are freed up so that pseudopods form and give rise to an irregular morphology. This focal adhesion turnover phenomenon also provides a rational justification for the 52% of prevalent “unconventional” cell motion and 20% of side protrusion, deciphered by the *CN correlation* analysis.

To understand how the cytoskeletal features impact cell polarity over the cell cycle, we also analyzed the distributions of FAs and SFs through their polar histograms in different cell cycle phases (**Fig. 4F**). After the polar histograms were determined through the rose plots with 10° frequency, the data show that both distributions are anisotropic in all cell cycle phases. The degree of isotropy can be further determined by the aspect ratio (AR) of the histograms, where a greater value indicates that the probed cytoskeletal features distributed with more bias, leading to a strong polarity. The results reveal the G1-phase cells have the highest AR value, followed by the G2-phase cells, then the S-phase cells with the lowest AR value. Thus, the polar histogram analysis reiterates that cells lose polarity when approaching the S phase, and regain some polarity in the G2 phase through focal adhesion turnover.

### Two CDKIs - p27^Kip1^ and p21^Cip1^ guide the RhoA and Rac1 signaling in migration evolutions

The effect of the cell cycle phases on cell motion suggests that there must be certain molecules involved in the underlying mechanism(s) between the cell cycle and cell dynamics. To our best knowledge, the only known proteins directly involved in these processes are two CDKIs, p27^Kip1^ and p21^Cip1^, which inhibit the CDKs in the nucleus of the G1-phase cells and RhoA and ROCK, respectively, in the cytoplasm after cells enter the S phase (Croft & Olson, 2006; Lee & Helfman, 2004). Thus, we documented the distributions of p27^Kip1^ and p21^Cip1^ in different cell cycle phases to observe whether their spatiotemporal appearance agrees with the activity profiles of RhoA and Rho kinase (ROCK) throughout the interphase of cell cycle.

First, we fluorescently labelled p27^Kip1^ and p21^Cip1^ in the synchronized NIH 3T3 fibroblasts of different cell cycle stages to quantitatively determine their cytoplasmic concentration profiles (**Fig. 5A**). The analysis shows that the cytoplasmic concentration of p27^Kip1^ continuously elevates throughout the cell cycle interphase, while cytoplasmic p21^Cip1^ exhibits a mild concave evolution with the lowest concentration appearing in the S phase (**Fig. 5B**). Consequently, we conducted quantitative fluorescence microscopy to determine the activity profiles of two Rho GTPases, RhoA and Rac1, that regulate membrane protrusion and actomyosin contraction, respectively (Olson *et al*, 1995) (**Fig. 5C**). The results show that the RhoA activity monotonically drops throughout the interphase of the cell cycle; whereas the Rac1 activity increases from the early G1 phase to the late G1 phase, it quickly declines to a minimum in the S phase, then elevates to a maximum in the G2 phase (**Fig. 5D**). The trends of these profiles are further supported through Western blot and qPCR techniques (**Fig. EV2**). Following these analyses, we also conducted active-ROCK pull down assay to determine the activity profile of ROCK in the interphase of the cell cycle (**Fig. 5E**).

**Figure 5.**
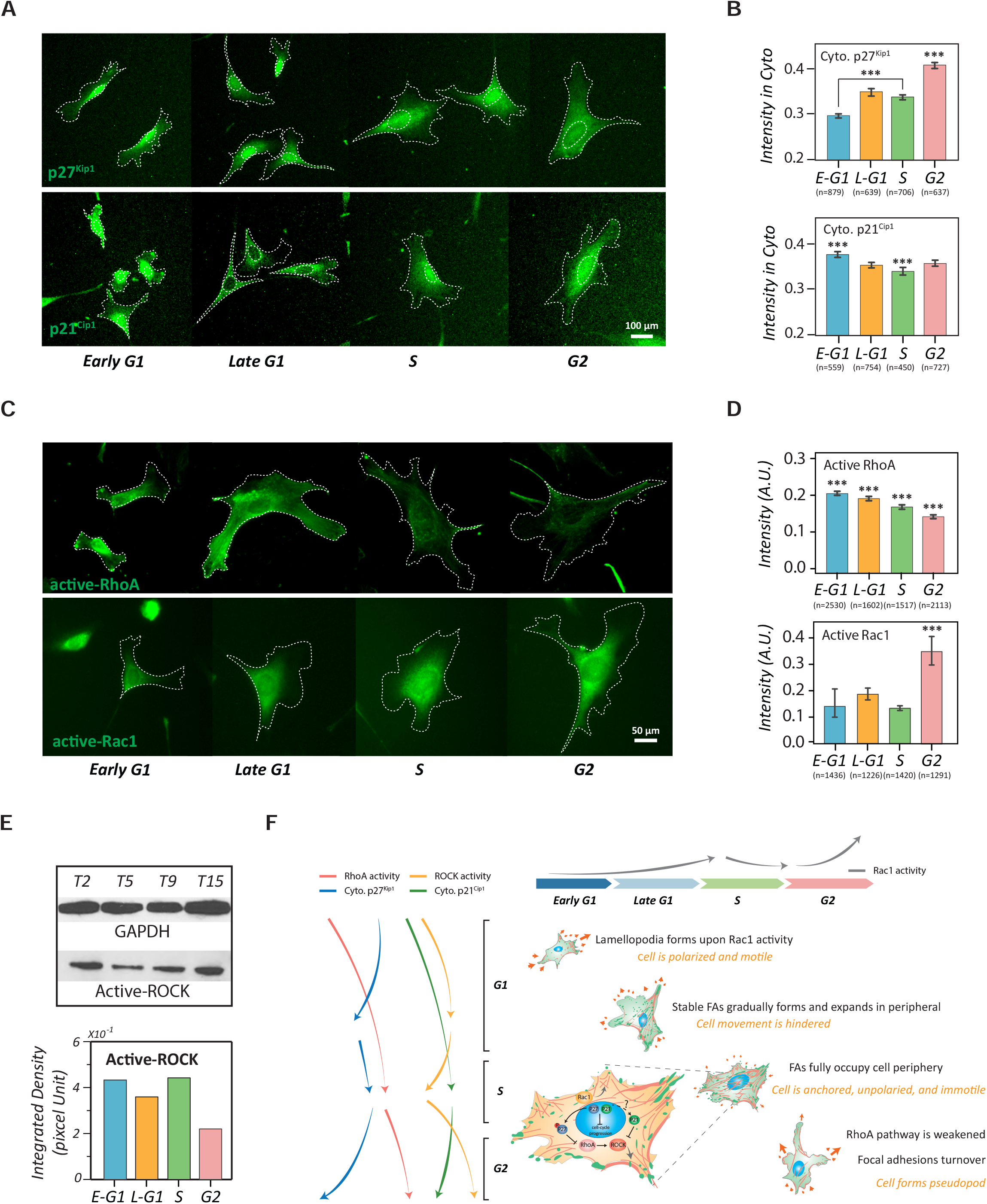
The involvement of cytoplasmic p27^Kip1^ and p21^Cip1^ in regulating the cell-cycle-dependent cell phenotype transitions. (**A**) Fluorescent images of p27^Kip1^ and p21^Cip1^ in cells of stages of the cell cycle. Dotted lines depict the boundaries of the cells and nuclei. Scale bar: 50 μm. (**B**) Bar plots showing the concentration profiles of the cytoplasmic p27^Kip1^ (up) and p21^Cip1^ (down) in different stages of the cell cycle phases. Error bars represent the SEM. (**C**) Immunostaining of active-RhoA and active-Rac1 in NIH 3T3 fibroblasts of each cell cycle stage. Dotted lines depict the boundaries of the cells. Scale bar: 50 μm. (**D**) Bar plots showing the activity profiles of active RhoA (up) and active Rac1 (down) in single cells of different cell cycle stages. (**E**) Western blot results of active ROCK in different cell cycle phases. Control: GAPDH. Error bars represent standard errors (n=2). (**F**) Metaphorical landscape illustrating the migration pattern transitions caused by Rho-GTPase and CDKIs pathways along with cell cycle progression. The magnified view of the S-phase cells shows the molecular interactions between CDKIs and RhoA pathways in the S phase. Data information: *** indicates that the group of data is statistically significant compared to other groups. The related statistical results of this figure are listed in Table EV4 and Table EV5.

The comparison of these profiles shows that a spatiotemporal agreement for the intermediate roles of p27^Kip1^ and p21^Cip1^ in the crossroad between the cell cycle and the cell movement. The increase of cytoplasmic p27^Kip1^ and p21^Cip1^ in cells from the S phase to the G2 phase is correlated with the decrease of RhoA and ROCK activities, respectively. Since the inhibition of RhoA pathway leads to the disassembly of focal adhesion (Kole *et al*, 2004), the weakening of RhoA pathway activity in the G2-phase cells supports the concurrence of focal adhesion turnover as we observed. In addition, the depleting of RhoA pathway activity only occurs after the p27^Kip1^ and p21^Cip1^ have been released from the CDK-cyclin complex (Besson *et al.*, 2004; Denicourt & Dowdy, 2004; Gui *et al.*, 2014; Lee & Helfman, 2004; Phillips *et al.*, 2018; Rampioni Vinciguerra *et al.*, 2019); hence, the RhoA pathway activity should be independent of the amounts of these two CDKIs in the G1-phase cells.

Immediately after cytokinesis, a newly divided cell enters the early G1 phase and spreads on the extracellular matrix (ECM). The cell explores its microenvironment via lamellipodia, which are formed upon the Rac1 activity. In this period, the cell is also polarized along the ECM that offers dense anchorage sites. Meanwhile, in the presence of mitogen, RhoA is activated, further promoting focal adhesions maturation in the polarized sites (Aplin & Juliano, 1999; Khosravi-Far *et al*, 1995; Olson *et al.*, 1995). The formation of dense, stable focal adhesions causes the hinderance of cell movement and the cell reaches the late G1 phase. During this time, the migration retention leads to prolonged substrate contacts around the cell periphery, inducing lateral protrusions through the activation of proximal Rac1 activity. When all the cell periphery is fully occupied by stable adhesions, the cell becomes stationary and enters the S phase. Starting around the same time, the presence of cytoplasmic CDKIs gradually diminishes the RhoA pathway activity, eventually causing focal adhesion turnover in the G2 phase. Hence, some parts of the cell periphery are freed up again and available for new cell-substrate contact developments, allowing the cell to form pseudopod-like features through re-surged Rac1 activity. This cytoskeletal remodeling alters cell morphology to be a form suitable for mitosis. Together, the evolution of cell phenotype is highly connected to the cell cycle progression (**Fig. 5F**).

## Discussion

This study reveals that both the morphology and dynamics of NIH 3T3 fibroblasts evolve along the interphase of the cell cycle with distinct molecular mechanisms. In the G1 phase, the most motile stage, the RhoA pathway plays an active role in promoting cell migration, while in the G2 phase, another dynamic interphase, the RhoA pathway becomes passive and CDKIs play a pivotal role in inhibiting the RhoA pathway and contributing to focal adhesion turnover to guide cytoskeletal remodeling. Furthermore, in addition to the existence of intratumor heterogeneity, cell-cycle-mediated drug resistance also allows some cancer cells to survive from the otherwise effective drugs and complicates prognostics(Beaumont *et al*, 2016; Shah & Schwartz, 2001; Yano *et al*, 2014). Hence, this study aims to avoid the confounding factors in heterogeneity caused by the cell cycle through suggesting a valuable strategy to evaluate the specific variables underlying drug resistance issues when applying target drugs in fighting cancers, making it necessary to take the molecular mechanisms and causes of heterogeneity into account. This study provides a framework to understand the molecular mechanisms that induce the heterogeneity of the cell phenotypes.

We made these solid conclusions based on three criteria: First, statistical evaluations reveal that all cell phenotypes, including motility, polarity, and morphology, maintain a high level of homogeneity in each stage of the cell cycle. The uniformity of these cellular behaviors in individual cell cycle phases is critical for the construction of the cellular/molecular framework. Second, the *CN correlation* approach allows us to decipher the cell dynamic patterns (*i.e.*, the quantitative contributions of different subcellular activities in the whole cell motion process) of single cells in different stages of the cell cycle. Finally, since the molecular mechanisms of individual subcellular activities in cell dynamics have been clearly determined (Kirfel *et al.*, 2004; Nobes & Hall, 1999; Olson *et al.*, 1995), the mechanisms of these subcellular activities have laid the foundations to thoroughly connect the activity profiles of the controlling molecules to the cell phenotype transitions over the cell cycle phases.

Despite the activity profiles of key proteins tightly correspond to a specific cell behavior, the whole cell physiology cannot be solely determined by discrete genomic and proteomic data due to the complex signaling crosstalk. However, the abundance of molecular information can be tightly integrated by single-cell behavior analysis platform, such as through the *CN correlation* analysis (Lan *et al.*, 2016; Lan *et al.*, 2018), which can collectively describe live-cell behaviors and their subcellular dynamic information in depth. A single-cell behavior analysis platform should be able to generate activity profiles from selected cell behavior with high sample sizes for statistical evaluations, therefore providing high-resolution information regarding the weighing contributions of composed subcellular activities. This capacity would allow the extension of applying machine deep learning into cell physiology.

## Materials and Methods

### Cell preparation

NIH 3T3 fibroblasts (American Type Culture Collection, Manassas, VA) were cultured in DMEM (Mediatech, Manassas, VA), supplemented with 10-% FBS (Hyclone Laboratories, Logan, UT), 1-% L-glutamine (Mediatech), and 1-% penicillin-streptomycin (Mediatech). Cells were maintained in an incubator with 10-% CO_2_ and at 37°C. The cell cultures were also kept below 70-% of confluency. For image acquisition, cell samples were prepared on the glass bottom dishes, which were pre-treated with 0.01-% poly-L-lysine (Sigma-Aldrich, St Louis, MO), following by 20-μl/ml fibronectin (BD Biosciences, San Jose, CA) coating.

### Cell synchronization

Cells were synchronized in the cell cycle using a double thymidine approach (Bostock *et al*, 1971). Two-mM thymidine (Sigma-Aldrich) was applied to the culture medium for 12 hours to arrest the cells at the stage of G1/S phase transition. Then, the cell culture was gently washed sequentially by HBSS (Mediatech), DMEM, and fresh culture medium, each for 15 minutes. Afterwards, cells were kept in regular culture medium for 9 hours before another 12-hour thymidine arrest. Finally, the cells were subjected to the same washing procedure described above to release the cells from cell cycle arrest. After the cells were released, the cell cycle profiles were determined hourly through individual cultured batches (see next section).

### Cell cycle profile determination using fluorescence microscopy

With the sample size of ~5000 cells, the nuclear intensity histogram was constructed using 50 bins and subjected to the Dean-Jett-Fox model (Dean & Jett, 1974; Fox, 1980) to determine the profile of the cell cycle phases. Briefly, normal curves with the same coefficient of variations were individually fitted to the G1 and the G2/M peaks of the histogram using least squares approximation. Then, the superposition of a normal curve and a broadened second-order polynomial was fitted to the histogram associated to the S phase. The areas covered by individual curves were used to determine the percentages of cells associated with the corresponding phases of the cell cycle.

### Cell cycle profile determination using flow cytometry

Individual batches of cell samples were detached from the culture dishes by trypsin and subjected to centrifugation. Then, the pellet was gently rinsed 3 times using ice-cold PBS before being fixed by 70-% ethanol overnight at 4°C. The next day, the pellet was re-suspended in PBS that contains 100-μg/ml ribonuclease, incubated for 30 minutes at room temperature, and subjected to 50-μg/ml propidium iodide (Abcam, Cambridge, MA) to label the DNA for 30 minutes at room temperature. Afterwards, LSR-II flow cytometer (BD Biosciences) were used to determine the percentage of cells in each phase of the cell cycle.

### Live-cell labelling

pEGFP plasmid (BD Biosciences) was transfected into NIH 3T3 fibroblasts using lipofectamine reagent (Invitrogen, Carlsbad, CA) based on a standard protocol. After transfection, the cells were cultivated onto the glass bottom dishes (World Precision Instrument, Sarasota, FL) by the single-cell density. After 24 hours, the cells were fluorescently labeled with GFP. Ten minutes before image acquisition, 20-μg/ml Hoechst 33342 was applied to the cell culture to label the nuclei (Siemann & Keng, 1986).

### Fixed cell labelling

Fibroblasts were fixed by 4-% paraformaldehyde (Sigma-Aldrich) in PBS for 15 minutes, permeabilized by 0.25-% Triton X-100 (Sigma-Aldrich) in PBS for 10 minutes, and blocked by 5-% BSA (Sigma-Aldrich) in PBS for 1 hour. For immunofluorescence staining, cells were incubated with primary antibody (listed in **Table EV6**) overnight at 4°C and washed 3 times by PBS. Then the corresponding fluorescently labeled secondary antibody was applied for 1 hour at room temperature before PBS washes. For F-actin and DNA staining, cells were incubated with 1:40 dilution of Alexa Fluor 568-phalloidin (Invitrogen) and 0.1-μg/ml Hoechst 33342. These stains were loaded with secondary antibodies at the same time.

### Microscopy and image acquisition

Fluorescent images were acquired using a TE-2000E imaging acquisition system (Nikon, Melville, NY), equipped with a 20× objective lens, a 60× oil-immersion objective lens, a digital eclipse C1 laser scanning confocal module, an X-Cite 120 PC fluorescent light source (EXFO, Ontario, Canada), and a Cascade: 1K CCD camera (Roper Scientific, Tucson, AZ). To avoid photobleaching, acquisition parameters were set at 300-ms exposure time, 3×3 binning, and 25-% maximal light intensity. All acquired signals were not saturated under such setting. During live-cell image acquisition, samples were placed in a CO_2_ supplementary system (In Vitro Scientific, St. Louis, MO), maintained at 10-% CO_2_ and 37 °C. Live-cell movies were recorded for 1 hour through two-channel fluorescence microscopy (the green channel for cells and the blue channel for nuclei) at 1-minute time intervals. Fixed fluorescent beads were bombarded on the glass-bottom slide during sample preparation to serve as the immobile references for image analysis to minimize positioning errors (Wu *et al.*, 2009; Wu *et al*, 2010).

### Rho GTPases pull-down assays

pGST-Rhotekin-RBD and pGST-PAK1-PBD plasmids (Addgene, Watertown, MA) were separately transformed into bacteria to produce the recombinant fusion proteins. Afterwards, glutathione Sepharose 4B resin (GE Healthcare Life Sciences, Pittsburg, PA) was added to the bacterial lysate for centrifugation. Then, the pellet was re-suspended in PBS with 10-% glycerol, stored at −80°C in 20-μl aliquot, and used as baits later for active RhoA and active Rac1 pull-down assay, respectively (Pellegrin & Mellor, 2008). Before pull-down, the aliquot was quantified using BSA as standard for consistency. Cell lysates were prepared in RIPA lysis buffer (10-mM Tris HCl, pH 7.4, 150-mM NaCl, 1-% (v/v) Triton X-100, 0.1-% (v/v) SDS, 0.5-% (v/v) DOC and 1-mM EDTA), containing protease inhibitor cocktail (Cytoskeleton, Inc., Denver, CO), and cleared by centrifugation (20,000 x g). Then, each lysate sample was mixed with an equal amount of pull-down resin and incubated at 4°C overnight. Next day, the sample was pulled down by centrifugation, washed 3 times by lysis buffer before the resin-bonded protein was dissociated by 2× SDS sample buffer for 2 minutes at room temperature. The sample was then boiled for 5 minutes, and subjected to 12-% SDS PAGE, following by Western blotting.

### Western blotting

After SDS-PAGE, the samples were transferred to the nitrocellulose membrane. Then, the membrane was blocked by 1X TBST with 5-% w/v nonfat dry milk for 1 hour at room temperature, and incubated with the primary antibody against the target protein overnight at 4°C. The next day, the membrane was washed with TBST, and incubated with horseradish-peroxidase-conjugated secondary antibody for 1 hour at room temperature. Detection was made by a standard protocol. The blotting results were quantified by ImageJ (NIH, Bethesda, MD). Antibodies are listed in **Table EV6**.

### Quantitative PCR to measure mRNA level

The total RNA in the cell sample was isolated by the RNeasy Mini Kit (Qiagen, Valencia, CA) and reverse-transcribed to cDNA by the iScript^TM^ Advanced cDNA Synthesis Kit (Bio-Rad, Hercules, CA). Then, the quantities of cDNA were determined through quantitative PCR (qPCR) using the CFX connect system (Bio-Rad). qPCR reactions were conducted using the SYBR supermix (Bio-Rad). The GAPDH quantities in cell samples were treated as the standard for cross-comparison. Intron-spanning primers for RhoA (JN971019.1), Rac1 (NM_009007.2), cyclin A1 (BC120518.1), cyclin E1 (BC138662.1), and GAPDH (GU214026.1) were designed using Primer3 and synthesized (Integrated DNA Technologies, Skokie, IL) as follows:

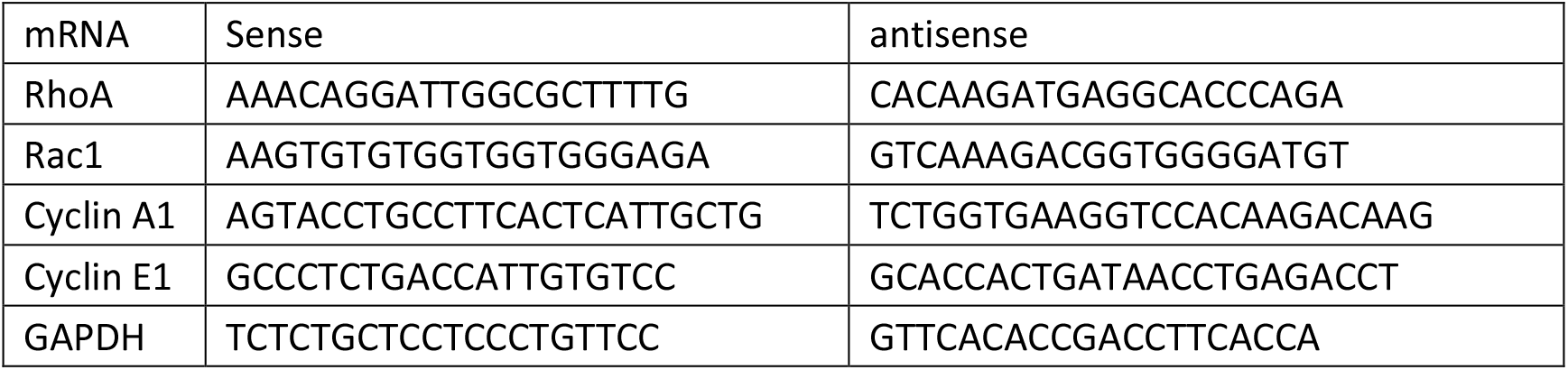

Measurements of each protein sample were conducted through 2 experimental duplicates and 2 biological duplicates. Relative cDNA levels were determined using the comparative Ct method (Schmittgen & Livak, 2008).

### Statistical Analysis

For live single-cell phenotype analysis, the sample sizes were greater than 20 cells. For fixed single-cell analysis, the sample sizes ranged from 200 to 2000. Data were displayed as mean ± SEM in either bar plot or box plot. The analysis of variance (ANOVA) was applied to compare the means of a condition among groups (more than 2 groups). When multiple group comparison by ANOVA showed significant differences exist among the compared group, the Tukey’s multi-comparison method would be applied to identify which groups are significantly different (*P* < *0.05*). The significant differences are labeled as *, **, and ***, to correspond to the *P* < *0.05*, *0.01*, and *0.001* criteria, respectively.

### The cell phenotype descriptors

The profiles of phenotype parameters were analyzed at 3-minute intervals (non-overlaid) over the cell cycle. Three significant cell shape phenotype parameters (Chen *et al*, 2016) were set as cell area, aspect ratio, and circularity: the area was identified by the actual number of pixels in the region times the scale unit per pixel (Arce *et al*, 2013); the aspect ratio defined as the length of the major axis divides by the length of the minor axis; where the cell has been normalized as the ellipse; the circularity value is computed as *circularity* = 4 × *Area* × π/*Perimeter*^2^.

### The partition of distinct data groups over the cell cycle progression

The profiles of instantaneous speeds and three shape parameters were initially randomly divided into three periods, then the least-squares criterion was applied among the data in the same parameters (groups) through an iterative process to obtain the final divisions:

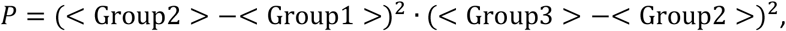

where each <> represents a mean data within the divided section.

### Homogeneity of cell long-term motility

To address the homogeneity of cell migration patterns within each cell cycle phase, we detected the convergence of *CMPI* value by continuously accumulating data by adding sample sizes until fulfilling the desired criterion (**Fig. EV2**). The analysis results reached a steady state (the standard error of the mean is less than 10% of the mean value) once the sample size exceeded 10. The information illustrated that cell migration is highly homogeneous and predictable when cells were synchronized to a specific cell cycle phase, thus supporting the claim that cell migration underlies cell cycle progression.

### SFs and FAs localization analysis

To quantify the distribution of SFs and FAs, ~50 cells were collected at a certain condition for SF/FA localization analysis in the individual-cell level. The intensity distributions of SFs and FAs in different cells were normalized and overlaid to a polar coordinate (with each cell centroid being set as origin, the orientation of the cell major axis being set as 0°, and a radius from each cell centroid to the radiated cell edges being set as 1) to describe the cell body (as seen in Fig. 4D). Then, *OpenAir* modules in *R* platform were applied to visualize the localization profiles occurrence frequencies of the SFs/FAs. The *polarPlot* function displayed a bivariate polar plot of SF/FA densities in different locations. Intensity were calculated by the local mean values (Fig. 4E). The *polarFreq* function displayed the distributions and occurrence frequencies of SFs/FAs in all directions, where the scale shows the occurrence counts in each bin (Fig. 4F).

**The computer vision approaches were performed by the MATLAB Image Processing toolbox (MathWorks, Natick, MA).** The key approaches are described as following:

### Cell and nuclear boundary determinations

A robust Gaussian fitting algorithms (Arce *et al.*, 2013) was applied to determine the cell boundaries. The nuclear boundaries were determined by direct segmentation using a given intensity threshold.

### Relative protein amount quantification

Under appropriate fluorescence microscopy setting (*i.e.*, intensity is not saturated), the amount of target protein was determined by the integrated greyscale value of the region of interest (*e.g.*, the cell, the cytoplasmic region, or the nucleus). The greyscale value was obtained from subtracting the background intensity by the mean intensity of the region of interest.

### The analysis of SFs and FAs

Signals of SFs were subjected to a sequence of processes from raw images: 1) Get rid of irrelevant background noise using reduced Gaussian filter (Arce *et al.*, 2013). 2) Adjust variance and magnitude of the signal using generic Gaussian and Laplacian filters. 3) Enhance crossing bright line using a linear Gaussian sensor (Eltzner *et al*, 2015). 4) Conduct local binarization via a combination of Gaussian weighted adaptive means and global thresholding (Eltzner *et al.*, 2015). 5) Segment SFs mask to get a skeleton of binary SFs mask. 6) Identify branch points in a skeleton structure by MATLAB built-in function *bwmorph*. 7) Identify different branches of fibers as the residual images by subtracting branch points from the skeleton. (8) Trace and reconnect these segmented fiber branches to enable reconstructed SFs (three basic criteria for connection procedure: the orientation of two compared fibers are less than 18°; the Euclidean maximum distance between two fiber branches endings are less than 8 pixels; and the mean intensity values and the width of fibers are to consistency, which is within 1.2 folds of difference) (**Fig. EV3A**).

Signals of FAs were subjected to a sequence of processes from raw images: 1) Get rid of irrelevant background noise using reduced Gaussian filter (Arce *et al.*, 2013). 2) Determine the local binarization by Otsu’s adjustable threshold method. 3) Apply morphological techniques called “opening-by-reconstruction” and “closing-by-reconstruction” to “clean” up the adjacent touching adhesion leagues. 4) Use a watershed segmentation approach to separate each FA (**Fig. EV3B**).

## Acknowledgements

We thank Professor Lizi Wu, Dr. Wee Ni, Dr. Qiangrong Yang, and Dr. Xin Zhou for technical consultants. We also thank Shihua Chen and Yeh Chen for their statistical analysis assistance. We thank Xinyi Xia for figure illustration. This work was partially supported by grants from NIH/NCI (U54CA143868) and DeBuYou Inc., China. All data is available to the public for reproducing purposes or further analysis.

## Author contributions

TL, YT, and JY conceived the study and designed experiments. TL, WC, HC, and JY conducted experiments and collected data. SW, LT, and ST designed and performed statistical data analysis. YT, TL, MY, SW, SS, and YZ discussed and wrote the manuscript.

## Expanded View Figure legends

**Fig. EV1.**
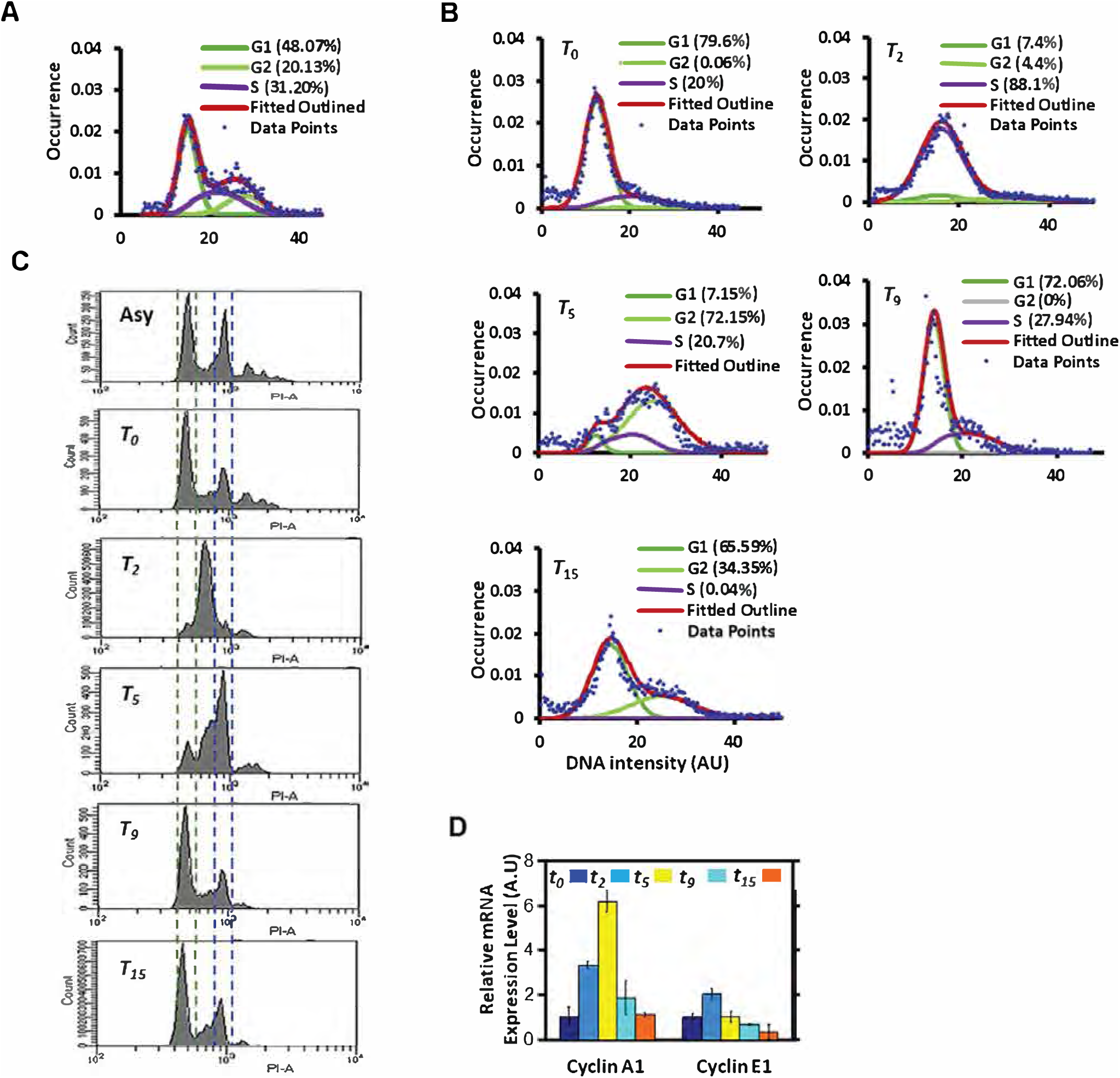
The cell cycle profile analyses of synchronized NIH 3T3 fibroblasts by time course, related to Fig. 1e. (**A**) The nuclear intensity profile of normal, unsynchronized cells. The profile (red) was subjected to the modified Dean-Jett fitting model to determine the weight of the cell population in each cell cycle phase: G1 (orange), S (purple), and G2 (green) phase of the cell cycle. (**B**) The nuclear intensity profiles and Dean-Jett fitting model of synchronized cells after released from the cell cycle arrest for a certain time course. The time of the optimal profiles of synchronized cells propagate to the G1/S, the early S, the S, the G2, the early G1, and the late G1 phase were determined as 0, 2, 5, 9, and 15 hours, respectively (and denoted as T0, T2, T5, T9, and T15, respectively). (**C**) The profiles of the cell cycle phases at the same time courses of cell propagation, after the cells were released from the cell cycle arrest, were verified by flow cytometry. (**D**) The amounts of cyclin A and cyclin E cDNAs, quantified at T0, T2, T5, T9, and T15 (left to right) to verify the cell cycle synchronization. Error bars represent the standard deviation (n=2).

**Fig. EV2.**
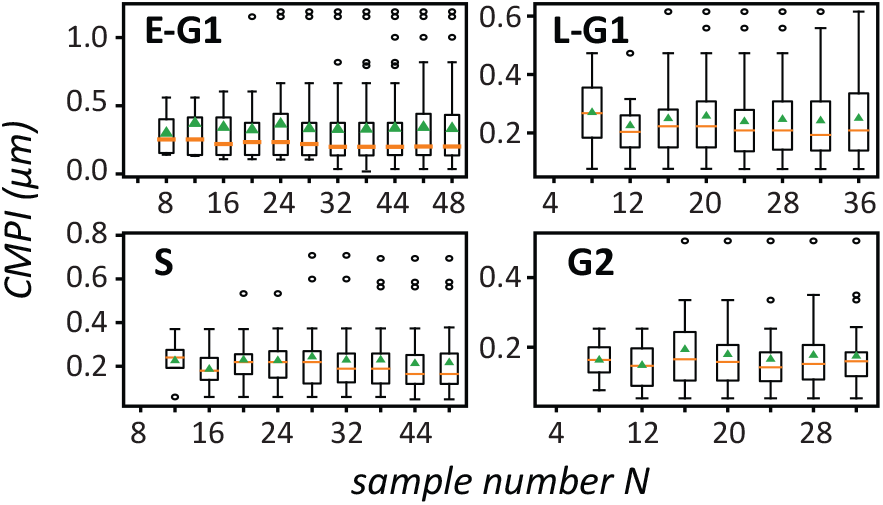
The convergence of long-term motility of cells in different cell cycle stages, estimated using CMPI. Box plots showing the convergences of *CMPI* against sample size in different stages of the cell cycle phases. Error bars represent standard error of the mean.

**Fig. EV3.**
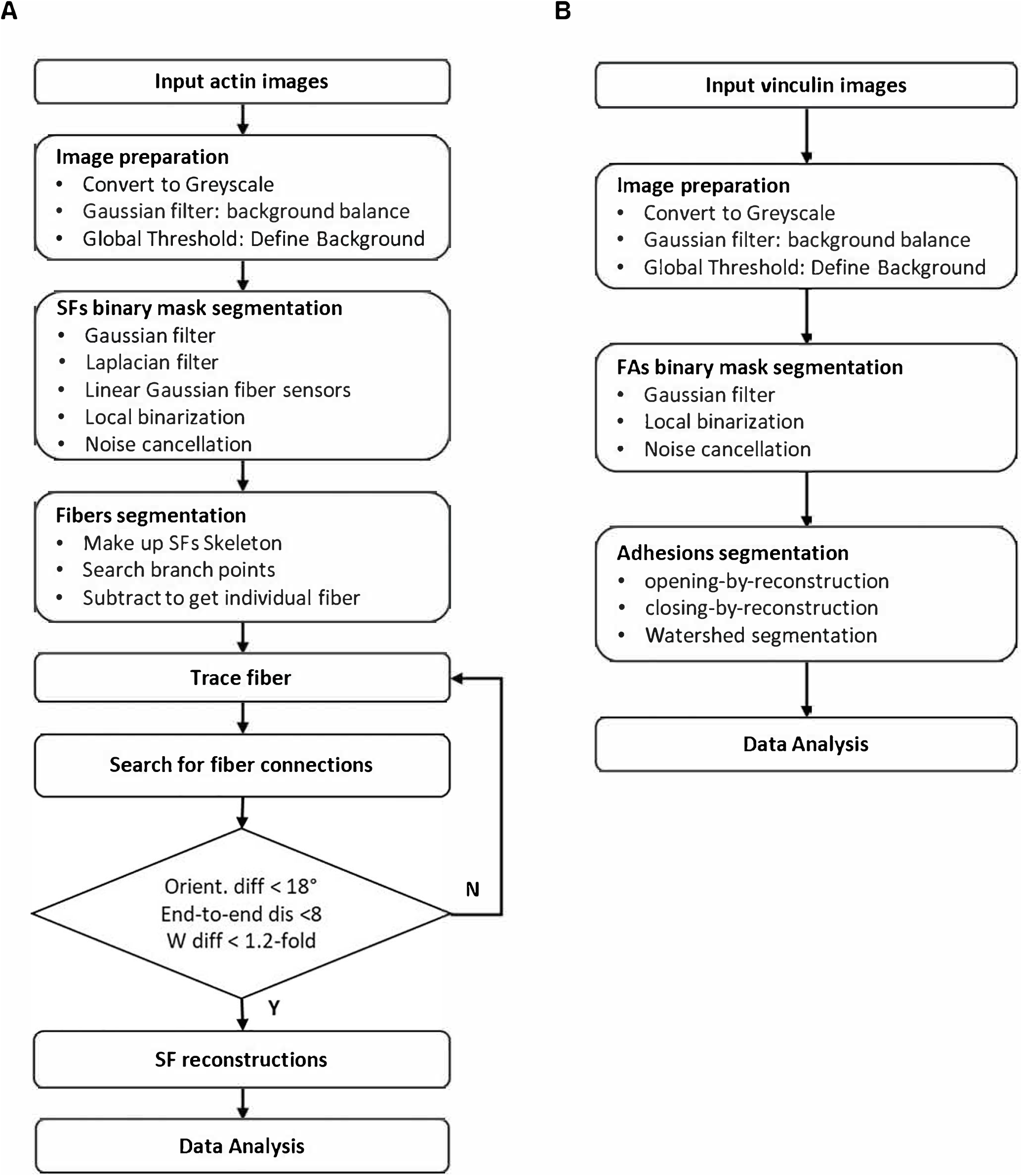
Flowcharts display the procedures of SF and FA analyses. (**A**) The image processing procedure of SFs. (**B**) The imaging processing procedure of FAs.

**Fig. EV4.**
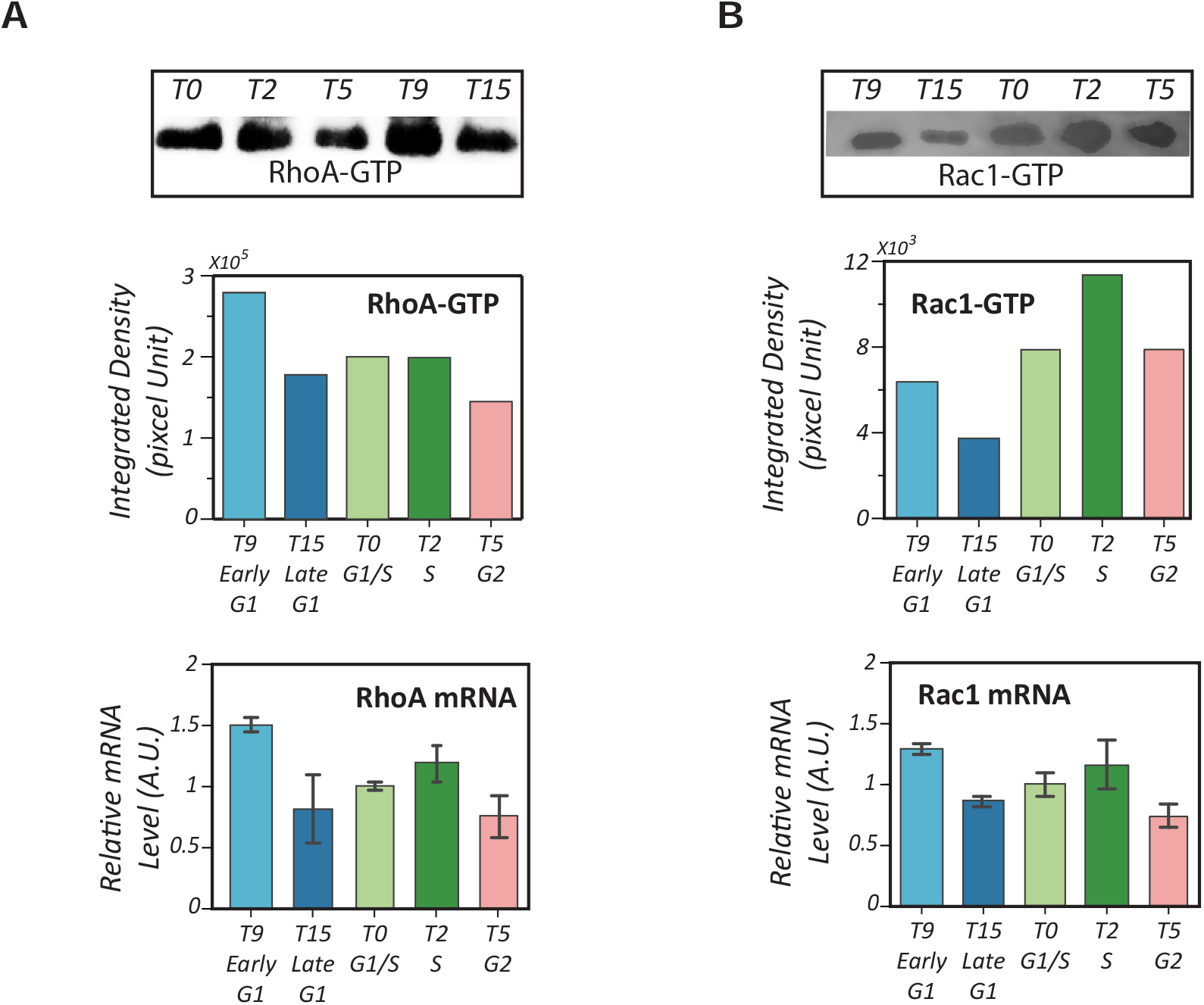
The activities and expression levels of RhoA and Rac1 in different cell cycle phases. Activities of RhoA and Rac1, assessed by RhoA-GTP and Rac1-GTP pull-down assays, respectively, followed by Western blots. ROCK activity was directly determined by Western blot using the Anti-phospho-MYPT1 (Thr696) antibody. (**A**) Western blotting results of the RhoA pull-down assay (upper panel) and the qPCR results of the total mRNA of RhoA (lower panel) in different stages of the cell cycle phases. (**B**) Western blotting results of the Rac1 pull-down assay (upper panel) and the qPCR results of the total mRNA of Rac1 (lower panel) in different stages of the cell cycle phases.

**Fig. EV5.**
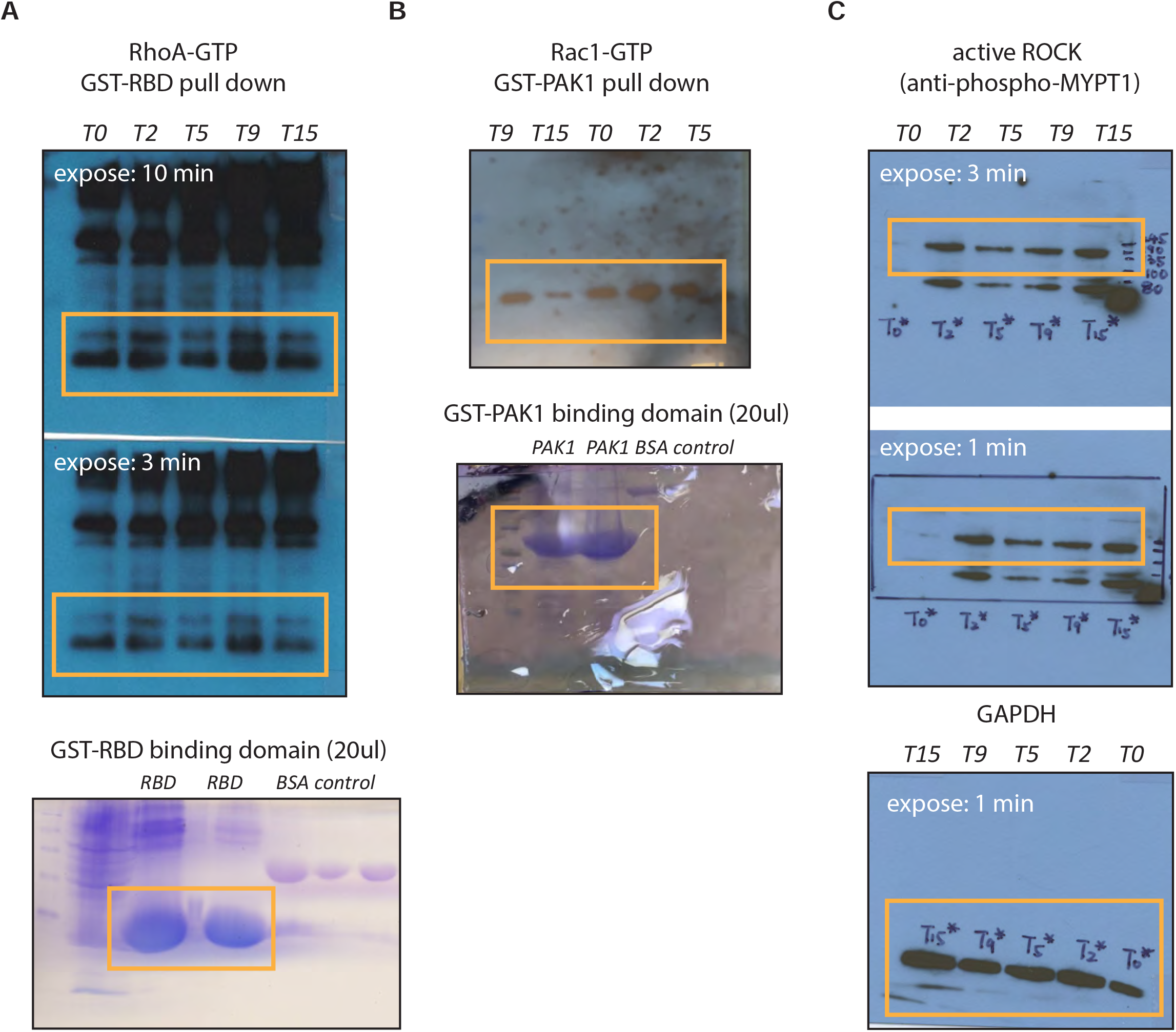
Original raw data related to Fig. EV4. (**A**) Original blotting results of RhoA-GTP pull-down assay and the associated SDS PAGE. (**B**) Original blotting results of Rac1-GTP pull-down assay and the associated SDS PAGE. (**C**) Western blotting results of active ROCK.

## Expanded View Tables and their legends

**Table EV1.** Statistical test results of Fig. 1F

| **NIH 3T3 fibroblast: Speed** |  |  |  |  |  |
| --- | --- | --- | --- | --- | --- |
| *One-way-ANOVA-Result (statistic=48.03, p-value=8.39e-31)* |  |  |  |  |  |
| *Multiple Comparison of Means - Tukey HSD, FWER=0.05* |  |  |  |  |  |
| **group1** | **group2** | **mean diff** | **lower** | **upper** | **reject** |
| *E-G1* | *G2* | -0.1845 | -0.2516 | -0.1174 | True |
| *E-G1* | *L-G1* | -0.0882 | -0.1529 | -0.0234 | True |
| *E-G1* | *S* | -0.2798 | -0.3427 | -0.217 | True |
| *G2* | *L-G1* | 0.0964 | 0.0241 | 0.1686 | True |
| *G2* | *S* | -0.0953 | -0.1659 | -0.0248 | True |
| *L-G1* | *S* | -0.1917 | -0.26 | -0.1233 | True |
| **NIH 3T3 fibroblast: Area** |  |  |  |  |  |
| *One-way-ANOVA-Result (statistic=472.8, p-value=2.0e-286)* |  |  |  |  |  |
| *Multiple Comparison of Means - Tukey HSD, FWER=0.05* |  |  |  |  |  |
| **group1** | **group2** | **mean diff** | **lower** | **upper** | **reject** |
| *E-G1* | *G2* | 1033.8488 | 931.5645 | 1136.1331 | True |
| *E-G1* | *L-G1* | 898.9655 | 800.2346 | 997.6964 | True |
| *E-G1* | *S* | 1113.4082 | 1017.6148 | 1209.2016 | True |
| *G2* | *L-G1* | -134.8833 | -245.0774 | -24.6893 | True |
| *G2* | *S* | 79.5594 | -28.0106 | 187.1294 | False |
| *L-G1* | *S* | 214.4427 | 110.2457 | 318.6398 | True |
| **NIH 3T3 fibroblast: Aspect Ratio** |  |  |  |  |  |
| *One-way-ANOVA-Result (statistic=101.76, p-value=7.71e-65)* |  |  |  |  |  |
| *Multiple Comparison of Means - Tukey HSD, FWER=0.05* |  |  |  |  |  |
| **group1** | **group2** | **mean diff** | **lower** | **upper** | **reject** |
| *E-G1* | *G2* | -0.6367 | -0.7996 | -0.4738 | True |
| *E-G1* | *L-G1* | -0.0165 | -0.1738 | 0.1407 | False |
| *E-G1* | *S* | -0.9037 | -1.0563 | -0.7511 | True |
| *G2* | *L-G1* | 0.6202 | 0.4447 | 0.7957 | True |
| *G2* | *S* | -0.267 | -0.4383 | -0.0957 | True |
| *L-G1* | *S* | -0.8872 | -1.0532 | -0.7213 | True |
| **NIH 3T3 fibroblast: Circularity** |  |  |  |  |  |
| *One-way-ANOVA-Result (statistic=44.3, p-value=2.02e-28)* |  |  |  |  |  |
| *Multiple Comparison of Means - Tukey HSD, FWER=0.05* |  |  |  |  |  |
| **group1** | **group2** | **mean diff** | **lower** | **upper** | **reject** |
| *E-G1* | *G2* | -0.0099 | -0.0234 | 0.0035 | False |
| *E-G1* | *L-G1* | -0.035 | -0.048 | -0.022 | True |
| *E-G1* | *S* | 0.0186 | 0.006 | 0.0312 | True |
| *G2* | *L-G1* | -0.025 | -0.0396 | -0.0105 | True |
| *G2* | *S* | 0.0285 | 0.0143 | 0.0427 | Ture |
| *L-G1* | *S* | 0.0535 | 0.0398 | 0.0673 | True |

**Table EV2.** Statistical test results of Fig. 3E

| **NIH 3T3 fibroblast: CMPI** |  |  |  |  |  |
| --- | --- | --- | --- | --- | --- |
| *One-way-ANOVA-Result (statistic=3.782, p-value=0.012)* |  |  |  |  |  |
| *Multiple Comparison of Means - Tukey HSD, FWER=0.05* |  |  |  |  |  |
| **group1** | **group2** | **mean diff** | **lower** | **upper** | **reject** |
| *E-G1* | *G2* | -0.1271 | -0.2432 | -0.011 | True |
| *E-G1* | *L-G1* | -0.0878 | -0.1998 | 0.0243 | False |
| *E-G1* | *S* | -0.1156 | -0.2243 | -0.0069 | True |
| *G2* | *L-G1* | 0.0393 | -0.0857 | 0.1644 | False |
| *G2* | *S* | 0.0115 | -0.1106 | 0.1336 | False |
| *L-G1* | *S* | -0.0278 | -0.1461 | 0.0904 | False |
| **NIH 3T3 fibroblast: 3min-CCD** |  |  |  |  |  |
| *One-way-ANOVA-Result (statistic=48.03, p-value=8.39e-31)* |  |  |  |  |  |
| *Multiple Comparison of Means - Tukey HSD, FWER=0.05* |  |  |  |  |  |
| **group1** | **group2** | **mean diff** | **lower** | **upper** | **reject** |
| *E-G1* | *G2* | -0.465 | -0.5768 | -0.3531 | True |
| *E-G1* | *L-G1* | -0.3794 | -0.4874 | -0.2715 | True |
| *E-G1* | *S* | -0.7796 | -0.8844 | -0.6749 | True |
| *G2* | *L-G1* | 0.0855 | -0.035 | 0.206 | False |
| *G2* | *S* | -0.3147 | -0.4323 | -0.197 | True |
| *L-G1* | *S* | -0.4002 | -0.5141 | -0.2863 | True |

**Table EV3.** Statistical test results of Fig. 4C

| **Focal adhesions (FAs): Number** |  |  |  |  |  |
| --- | --- | --- | --- | --- | --- |
| *One-way-ANOVA-Result (statistic=3.82, p-value=0.025)* |  |  |  |  |  |
| *Multiple Comparison of Means - Tukey HSD, FWER=0.05* |  |  |  |  |  |
| group1 | group2 | mean diff | lower | upper | reject |
| E-G1 | G2 | -28.7037 | -77.5016 | 20.0942 | False |
| E-G1 | S | -55.4259 | -103.3848 | -7.4671 | True |
| G2 | S | -26.7222 | -80.0255 | 26.5811 | False |
| **Focal adhesions (FAs): Area** |  |  |  |  |  |
| *One-way-ANOVA-Result (statistic=91.09, p-value=3.84e-40)* |  |  |  |  |  |
| *Multiple Comparison of Means - Tukey HSD, FWER=0.05* |  |  |  |  |  |
| group1 | group2 | mean diff | lower | upper | reject |
| E-G1 | G2 | 2.435 | 1.7183 | 3.1518 | True |
| E-G1 | S | -1.5732 | -2.2357 | -0.9107 | True |
| G2 | S | -4.0083 | -4.7047 | -3.3118 | Ture |
| **Focal adhesions (FAs): Density** |  |  |  |  |  |
| *One-way-ANOVA-Result (statistic=1.92, p-value=0.151)* |  |  |  |  |  |
| *Multiple Comparison of Means - Tukey HSD, FWER=0.05* |  |  |  |  |  |
| group1 | group2 | mean diff | lower | upper | reject |
| E-G1 | G2 | -0.0069 | -0.021 | 0.0071 | False |
| E-G1 | S | 0.0036 | -0.009 | 0.0163 | False |
| G2 | S | 0.0106 | -0.0023 | 0.0234 | False |
| **Stress Fibers (SFs): Number** |  |  |  |  |  |
| *One-way-ANOVA-Result (statistic=10.95, p-value=4.25e-05)* |  |  |  |  |  |
| *Multiple Comparison of Means - Tukey HSD, FWER=0.05* |  |  |  |  |  |
| group1 | group2 | mean diff | lower | upper | reject |
| E-G1 | G2 | -118.4368 | -198.3764 | -38.4973 | True |
| E-G1 | S | -139.7685 | -218.3336 | -61.2034 | True |
| G2 | S | -21.3317 | -108.652 | 65.9886 | False |
| **Stress Fibers (SFs): Length** |  |  |  |  |  |
| *One-way-ANOVA-Result (statistic=66.52, p-value=1.44e-29)* |  |  |  |  |  |
| *Multiple Comparison of Means - Tukey HSD, FWER=0.05* |  |  |  |  |  |
| group1 | group2 | mean diff | lower | upper | reject |
| E-G1 | G2 | -2.0542 | -2.4536 | -1.6547 | True |
| E-G1 | S | -0.4383 | -0.8266 | -0.0501 | True |
| G2 | S | 1.6158 | 1.2189 | 2.0127 | Ture |
| **Stress Fibers (SFs): Density** |  |  |  |  |  |
| *One-way-ANOVA-Result (statistic=0.84, p-value=0.43)* |  |  |  |  |  |
| *Multiple Comparison of Means - Tukey HSD, FWER=0.05* |  |  |  |  |  |
| group1 | group2 | mean diff | lower | upper | reject |
| E-G1 | G2 | 0.0089 | -0.0085 | 0.0263 | False |
| *E-G1* | *S* | 0.0065 | -0.0106 | 0.0236 | False |
| *G2* | *S* | -0.0024 | -0.0214 | 0.0166 | False |

**Table EV4.** Statistical test results of Fig. 5B

| **NIH 3T3 fibroblast: P27Kip1** |  |  |  |  |  |
| --- | --- | --- | --- | --- | --- |
| *One-way-ANOVA-Result (statistic=76.68, p-value=1.07e-47)* |  |  |  |  |  |
| *Multiple Comparison of Means - Tukey HSD, FWER=0.05* |  |  |  |  |  |
| **group1** | **group2** | **mean diff** | **lower** | **upper** | **reject** |
| *E-G1* | *G2* | -0.0569 | -0.0693 | -0.0445 | True |
| *E-G1* | *L-G1* | 0.0024 | -0.01 | 0.0148 | False |
| *E-G1* | *S* | 0.0139 | 0.0019 | 0.026 | True |
| *G2* | *L-G1* | 0.0593 | 0.0459 | 0.0727 | True |
| *G2* | *S* | 0.0708 | 0.0578 | 0.0839 | True |
| *L-G1* | *S* | -0.0115 | -0.0246 | 0.0015 | False |
| **NIH 3T3 fibroblast: P27Kip1** |  |  |  |  |  |
| *One-way-ANOVA-Result (statistic=76.68, p-value=1.07e-47)* |  |  |  |  |  |
| *Multiple Comparison of Means - Tukey HSD, FWER=0.05* |  |  |  |  |  |
| **group1** | **group2** | **mean diff** | **lower** | **upper** | **reject** |
| *E-G1* | *G2* | -0.0569 | -0.0693 | -0.0445 | True |
| *E-G1* | *L-G1* | 0.0024 | -0.01 | 0.0148 | False |
| *E-G1* | *S* | 0.0139 | 0.0019 | 0.026 | True |
| *G2* | *L-G1* | 0.0593 | 0.0459 | 0.0727 | True |
| *G2* | *S* | 0.0708 | 0.0578 | 0.0839 | True |
| *L-G1* | *S* | -0.0115 | -0.0246 | 0.0015 | False |

**Table EV5.** Statistical test results of Fig. 5D

| **NIH 3T3 fibroblast: active-RhoA** |  |  |  |  |  |
| --- | --- | --- | --- | --- | --- |
| *One-way-ANOVA-Result (statistic=110.78, p-value=3.11e-70)* |  |  |  |  |  |
| *Multiple Comparison of Means - Tukey HSD, FWER=0.05* |  |  |  |  |  |
| **group1** | **group2** | **mean diff** | **lower** | **upper** | **reject** |
| *E-G1* | *G2* | 0.0367 | 0.0313 | 0.0421 | True |
| *E-G1* | *L-G1* | 0.0083 | 0.0025 | 0.0142 | True |
| *E-G1* | *S* | 0.0214 | 0.0154 | 0.0273 | True |
| *G2* | *L-G1* | -0.0284 | -0.0345 | -0.0223 | True |
| *G2* | *S* | -0.0153 | -0.0215 | -0.0091 | True |
| *L-G1* | *S* | -0.0131 | -0.0196 | -0.0065 | True |
| **NIH 3T3 fibroblast: active-Rac1** |  |  |  |  |  |
| *One-way-ANOVA-Result (statistic=22.04, p-value=3.50e-14)* |  |  |  |  |  |
| *Multiple Comparison of Means - Tukey HSD, FWER=0.05* |  |  |  |  |  |
| **group1** | **group2** | **mean diff** | **lower** | **upper** | **reject** |
| *E-G1* | *G2* | -0.2057 | -0.2815 | -0.1299 | True |
| *E-G1* | *L-G1* | -0.0453 | -0.1222 | 0.0316 | False |
| *E-G1* | *S* | 0.0068 | -0.0672 | 0.0808 | False |
| *G2* | *L-G1* | 0.1604 | 0.0815 | 0.2392 | True |
| *G2* | *S* | 0.2125 | 0.1365 | 0.2886 | True |
| *L-G1* | *S* | -0.0521 | -0.1292 | 0.025 | False |

**Table EV6.**
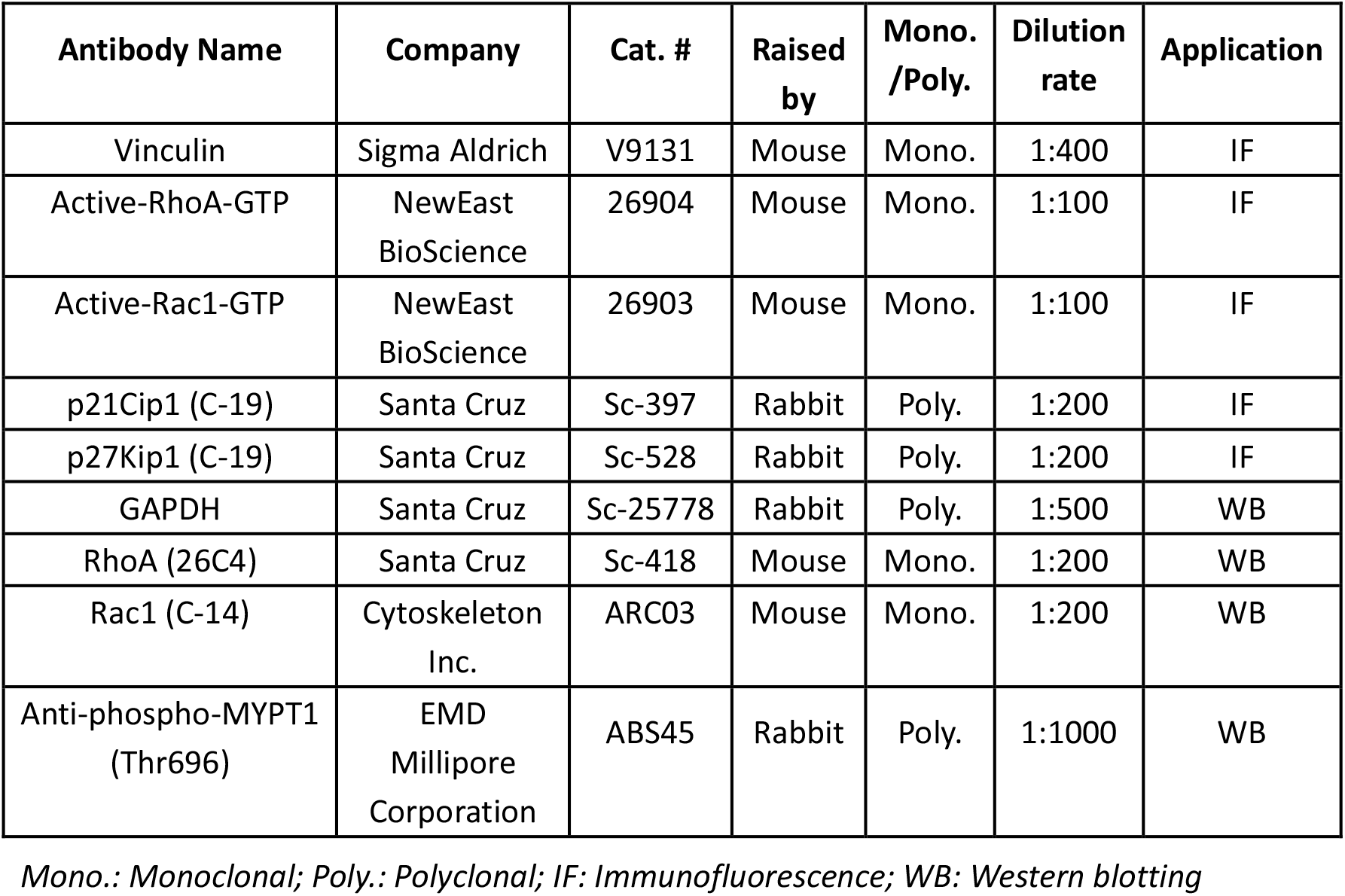
Primary antibodies used in this study.

